# Modular Deep Learning for Direct RNA Sequence Design via Self-Contained RNA Units

**DOI:** 10.64898/2026.04.16.719021

**Authors:** Jian Wang, Nikolay V. Dokholyan

## Abstract

RNA sequence design is a pivotal challenge in synthetic biology, yet state-of-the-art deep learning methods face a fundamental bottleneck: the scarcity of high-resolution 3D structures. To compensate for limited training data, existing approaches like NA-MPNN and RiboDiffusion employ computationally expensive autoregressive or iterative diffusion sampling, substantially limiting their throughput and scalability. In this work, we propose that this data limitation is largely a problem of accessibility and granularity. We introduce SCRU-DB, a comprehensive database that systematically decomposes complex RNAs into over 61,000 Self-contained RNA Units (SCRUs). This scale far exceeds previous RNA motif libraries, capturing over 8,200 unique structural clusters. Crucially, SCRUs are rigorously defined as structurally autonomous modules identified via tertiary contact clustering, ensuring they act as self-stabilizing, foldable physical units. Leveraging this massive, modular prior, we present SCRU-Seq (a direct, O(1) prediction GNN) and SCRU-Diff (an iterative diffusion model). On our high-fidelity set112 benchmark, SCRU-Seq achieves a native sequence recovery (NSR) of 63.7%, while SCRU-Diff reaches a superior Best NSR of 79.2%. We demonstrate high structural fidelity via 3D backbone superposition using the C4’ RMSD (reaching 1.5Å for complex targets) and validate the structural isomorphism of our modular fragments. This framework provides a scalable, physically grounded solution for generating diverse and structurally accurate RNA sequences.

## Introduction

Ribonucleic acid (RNA) has transcended its traditional role as a passive genetic messenger to become a central pillar of modern therapeutics and synthetic biology. From mRNA vaccines^1^ and CRISPR-Cas9^2^ guide RNAs to catalytic ribozymes and sensing riboswitches, the function of these molecules is strictly dictated by their ability to fold into precise three-dimensional (3D) architectures. Consequently, RNA design^3–5^, the inverse problem of identifying a nucleotide sequence that will stably adopt a target backbone structure, has emerged as a critical computational challenge. Solving this problem allows us to engineer novel molecular machines with specific affinities and catalytic activities, accelerating drug discovery and the development of programmable genetic circuits.

Historically, the majority of RNA design efforts have utilized secondary structure (2D) abstractions^5–14^, focusing on sequences that satisfy specific base-pairing rules. However, 2D-centric design is fundamentally limited as it fails to capture the intricate tertiary interactions, including non-canonical base pairs and long-range spatial packing, that define the functional state of natural RNAs. These 3D architectural features are indispensable for the formation of catalytic centers and precise binding pockets, which are invisible to discrete 2D topologies. Furthermore, designing directly in 3D space allows for the explicit consideration of continuous backbone geometry and global steric constraints, ensuring that the designed sequences are not merely "pairing-compatible" but are physically capable of folding into the intended, compact global architecture.

Current computational approaches to RNA design generally fall into two categories: physics-based optimization and deep learning. Traditional physics-based methods, such as RNAinverse or Rosetta’s rna_denovo, rely on stochastic search algorithms to minimize free energy. While thermodynamically rigorous, these methods are computationally intensive and often struggle to traverse the rugged energy landscapes of large RNAs, limiting their utility for high-throughput applications.

Recently, deep learning methods have shown great promise by learning geometric constraints directly from structural data. Prominent examples include NA-MPNN^15^, a message-passing graph neural network that treats nucleic acids and proteins within a unified framework, and RiboDiffusion^4^, a generative diffusion model that iteratively refines sequences. NA-MPNN employs an autoregressive decoder, generating sequences one nucleotide at a time (𝑂(𝐿) complexity), while RiboDiffusion relies on a denoising process requiring hundreds of iterative steps (𝑇 steps). Although these sophisticated sampling strategies, such as Autoregressive (AR) generation and Diffusion, are designed to capture complex probability distributions, we argue that they are essentially compensatory mechanisms for a more fundamental limitation: data scarcity. The number of solved high-resolution RNA structures in the PDB is orders of magnitude smaller than that of proteins. By training on full-length global structures, these models effectively starve for data, forcing reliance on computationally expensive inference procedures to squeeze out marginal performance gains from a limited training set.

Our core hypothesis is that the bottleneck in RNA design is not the complexity of the generative model, but the richness of the structural priors it learns from. We observe that the perceived “data scarcity” is largely a problem of accessibility and granularity. The RNA structural landscape is strictly dominated by massive macromolecular complexes, such as the Ribosome, where individual chains frequently exceed thousands of nucleotides. These colossal structures are built from repeating, modular substructures. By systematically decomposing these large RNAs into smaller, learnable units, we can effectively unlock the vast structural information hidden within them. This process converts a limited set of global PDB entries (9,406 structures as of March 2026) into a massive, high-density dataset of 61,916 training examples, a nearly 7-fold expansion in available data. This scale far exceeds previous motif libraries, capturing over 8,200 unique structural clusters spanning wide length and complexity spectra.

However, the method of decomposition is critical. We posit that standard secondary structure elements (SSEs), as traditionally defined in hierarchical folding (e.g., RNAComposer^16^, MC-Sym^17^, and 3dRNA^18,19^), are fundamentally unsuitable for training design models. A bare SSE, such as an internal loop, is often thermodynamically unstable in isolation. Using such fragments introduces critical noise, creating a physically invalid sequence-structure mapping. To overcome this, we propose SCRU-DB, a comprehensive database built on the concept of the <u>S</u>elf-<u>C</u>ontained <u>R</u>NA <u>U</u>nit (SCRU). Rather than relying on isolated motifs or hierarchical tree abstractions, a SCRU is constructed as a self-contained unit by selectively combining multiple helical regions, which provide the foundational structural stability through dense base-pairing, with the fragments that navigate between them. By representing the RNA structure as a connectivity graph rather than a nested tree, our framework inherently captures the topological complexity of the folding landscape and provides natural support for pseudoknots. This ensures that each SCRU serves as a thermodynamically stable physical unit that exhibits structural isomorphism, retaining its precise native fold regardless of whether it is in isolation or part of a global chain. By decomposing PDB structures into these robust units, we generate a training set that explicitly captures learnable, high-fidelity physicochemical constraints.

Leveraging this data-centric breakthrough, we introduce two generative models built upon the SCRU-DB: SCRU-Seq, a direct-prediction GNN that provides high-speed inference (𝑂(1)), and SCRU-Diff, a diffusion model designed to capture the one-to-many nature of the RNA design problem. Both models employ a unique Dual-Radius Graph architecture that simultaneously captures dense local stereo-chemistry (via all-atom connections < 4Å) and sparse global topology (via C4’ backbone connections < 20Å). On our benchmark consisting of 112 RNAs, SCRU-Seq achieves a native sequence recovery (NSR) of 63.7%, while SCRU-Diff pushes the generative potential further, reaching a Best NSR of 79.2%. We find that our modular training approach allows for the generation of sequences that accurately adopt native 3D geometries, with C4’ RMSD values as low as 1.5Å. This framework validates our premise that with the right modular representation (SCRU-DB), RNA design can be solved effectively and scalably.

## Results

### Overview of the Modular RNA Design Workflow

We introduce a modular RNA design framework that addresses data scarcity through a systematic deconstruction-reconstruction pipeline (Figure 1). We begin by decomposing high-resolution RNA 3D structures from the RCSB PDB (Figure 1a) into a connectivity graph representation, where stable helical regions, the primary anchors of structural stability, and their intervening fragments are identified (Figure 1b). This graph-based representation marks a fundamental departure from traditional hierarchical models, such as the RNA secondary structure tree. In our framework, nodes specifically represent continuous helical regions defined by canonical base pairing, while edges represent the flexible linking fragments that traverse between these anchors. Unlike a hierarchical tree, which is restricted to nested parent-child relationships and often fails to represent non-nested interactions, our connectivity graph allows for arbitrary topological connections. By enabling cyclic dependencies and cross-linking between distal helical nodes, this architecture naturally accommodates pseudoknots and long-range tertiary interactions that are typically excluded or simplified in tree-based methods. This ensures that each extracted SCRU captures the complete topological and thermodynamic constraints of the molecule, serving as a physically autonomous module capable of folding independently.

**Figure 1:**
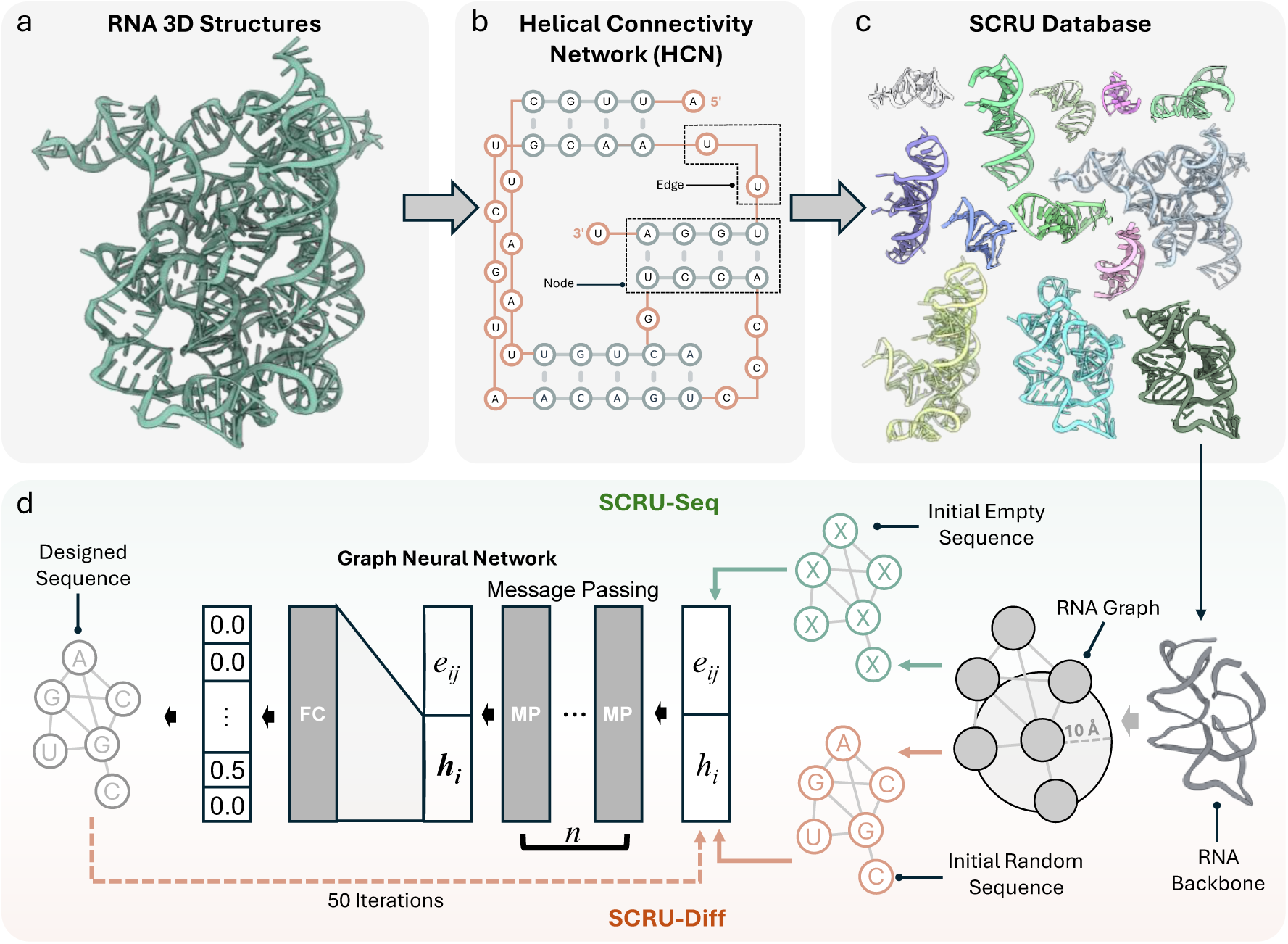
SCRU-Seq Design Workflow. Overview of the modular RNA design pipeline. - a: Representative large-scale RNA 3D structures from the PDB. - b: Hierarchical decomposition of complex RNA global folds into a Generalized Hairpin Tree (GHairpin tree). - c: The resulting SCRU Database (SCRU-DB) containing modular units (Stems, Hairpins, Junctions) extracted for scalable training. - d: Model architectures: SCRU-Seq (GNN-based direct prediction) and SCRU-Diff (iterative refinement process). Both models utilize dual-radius bipartite graphs that integrate dense local atomic environments with sparse global topology.

We leverage the resulting SCRU Database (SCRU-DB) (Figure 1c) as a high-density training resource. Based on this database, we developed two parallel generative models (Figure 1d): SCRU-Seq, a dual-radius bipartite graph neural network for high-speed direct prediction, and SCRU-Diff, an iterative diffusion-based model designed for high-diversity sequence generation. During our benchmarking, we observed that while both models achieve comparable design accuracy, SCRU-Diff explores a broader sequence space, offering unique advantages for diverse candidate exploration.

### Quantitative Characterization of the SCRU-DB

Through a multi-dimensional analysis of the SCRU-DB, we reveal the scale and complexity of the structural landscape captured by our partitioning algorithm (Figure 2). We identify SCRUs by modeling RNA as a connectivity graph of helical regions, which provide the foundational thermodynamic stability through dense base-pairing, linked by intervening sequence fragments (Figure 2a). Critically, we differentiate SCRUs from traditional secondary structure motifs (e.g., standard hairpins, internal loops, or junctions found in existing libraries). While traditional motifs often isolate the central loop, our SCRU definition identifies structurally autonomous modules by clustering secondary structure elements based on tertiary interactions and the length of connecting helical bridges. By treating the RNA as a graph rather than a nested tree, nuestra approach captures topological complexities and naturally accommodates pseudoknots, ensuring that SCRUs are thermodynamically stable building blocks that retain their native fold in isolation—a property we define as structural isomorphism.

**Figure 2:**
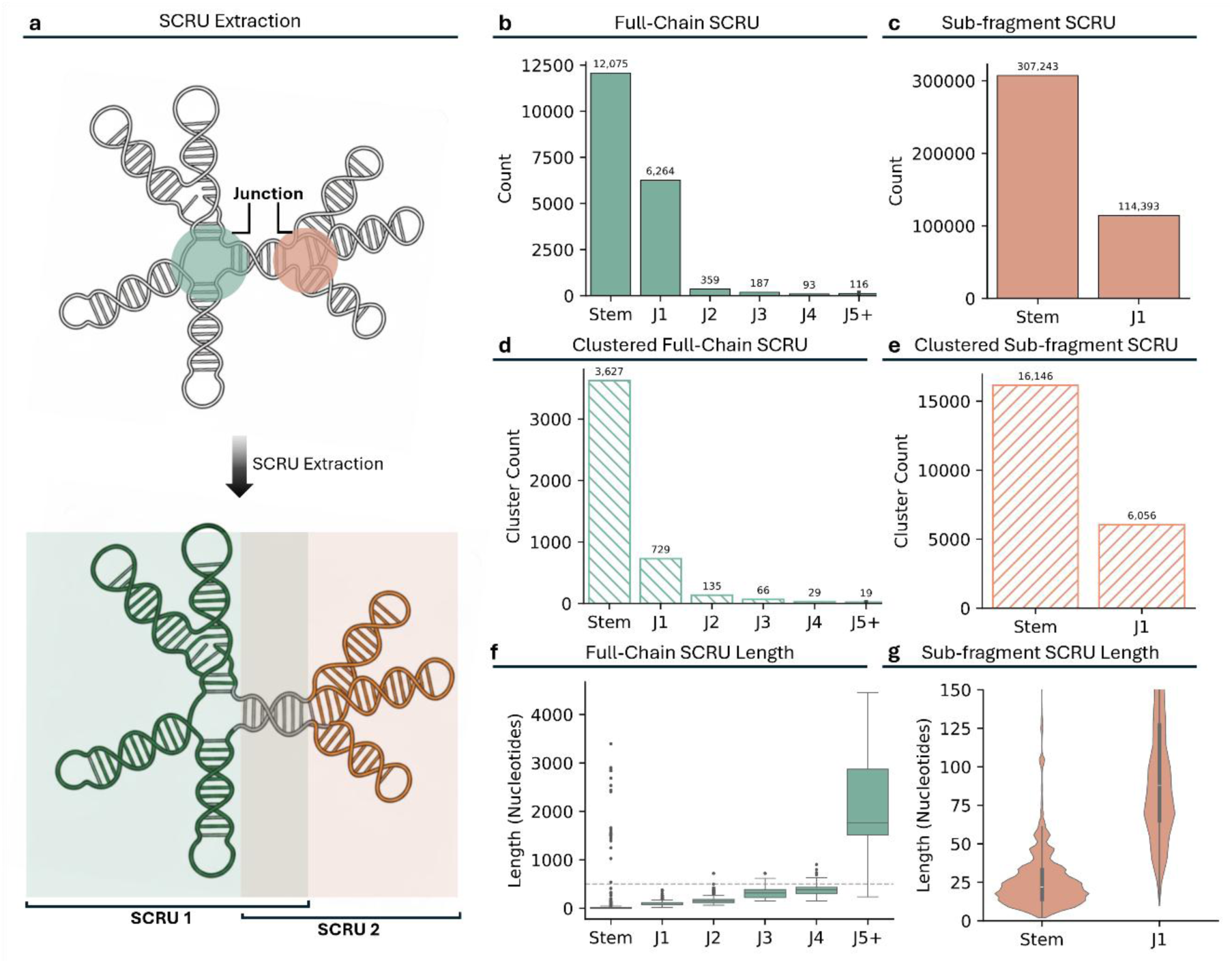
SCRU-DB Statistics and Characterization. Multi-dimensional analysis of the Smallest Consecutive RNA Units (SCRU) database. - a: Schematic of the topological decomposition used to identify Junctions and Stems. - b, c: Distribution of SCRU types for Full-Chain (b) and Sub-fragment (c) datasets. - d, e: Number of distinct structural clusters identified by MMseqs2 for Full-Chain (d) and Sub-fragment (e) categories. - f: Nucleotide length distribution of Full-Chain SCRUs, categorized by junction complexity (J1 to J5+, representing the number of stems connected at a junction). - g: Sequence length distribution of Sub-fragment SCRUs (Violin plot), highlighting the high density of structural motifs within the database.

Our database provides a massive expansion over previous structural resources. While benchmarks like the BGSU RNA 3D Motif Atlas^20^ typically contain approximately 15,000-25,000 total motif instances across ∼3,000 unique groups, and the RNA-As-Graphs Motif Atlas^21^ identifies around 1,844 distinct submotifs, SCRU-DB converts 9,406 high-resolution PDB entries into over 61,916 modular SCRU units. This represents a nearly 7-fold increase in available training data compared to global PDB entries and more than an order of magnitude increase in instance volume over specialized libraries. To ensure diversity, we identified approximately 8,405 unique sequence-structure clusters (4,386 sub-fragment and 4,019 full-chain) via MMseqs2 (Figure 2d, e), providing a significantly larger and more balanced generative space than any existing structural dataset.

We categorize the extracted units based on their topological complexity (junction complexity). In this classification, J1 refers to simple single-stem hairpins, J2 to 2-way junctions such as internal loops or bulges, and J3 through J5+ to complex multi-way branching hubs where three or more helical arms originate from a central loop (Figure 2b). Our database maintains robust representation across this entire spectrum, from simple J1 hairpins to massive J5+ ribosomal hubs, providing the most diverse modular RNA training set to date. Our length analysis (Figure 2f, g) further highlights this dual-scale nature, spanning small modular fragments (median < 25 nt) to massive complexes exceeding 4,000 nucleotides.

### Native Sequence Recovery Performance Benchmarking

We evaluated both design models on a non-redundant independent benchmark of full-length RNA chains. To ensure a rigorous and fair comparison, we curated this benchmark through a multi-stage filtering pipeline (Figure 3a). First, we performed a global cluster-level split of the SCRU-DB using MMseqs2 (80% identity threshold), assigning all structural motifs from the same sequence cluster to either the training or test split to prevent data leakage. This yielded an initial pool of 2,129 full-chain RNAs in the independent test split.

**Figure 3:**
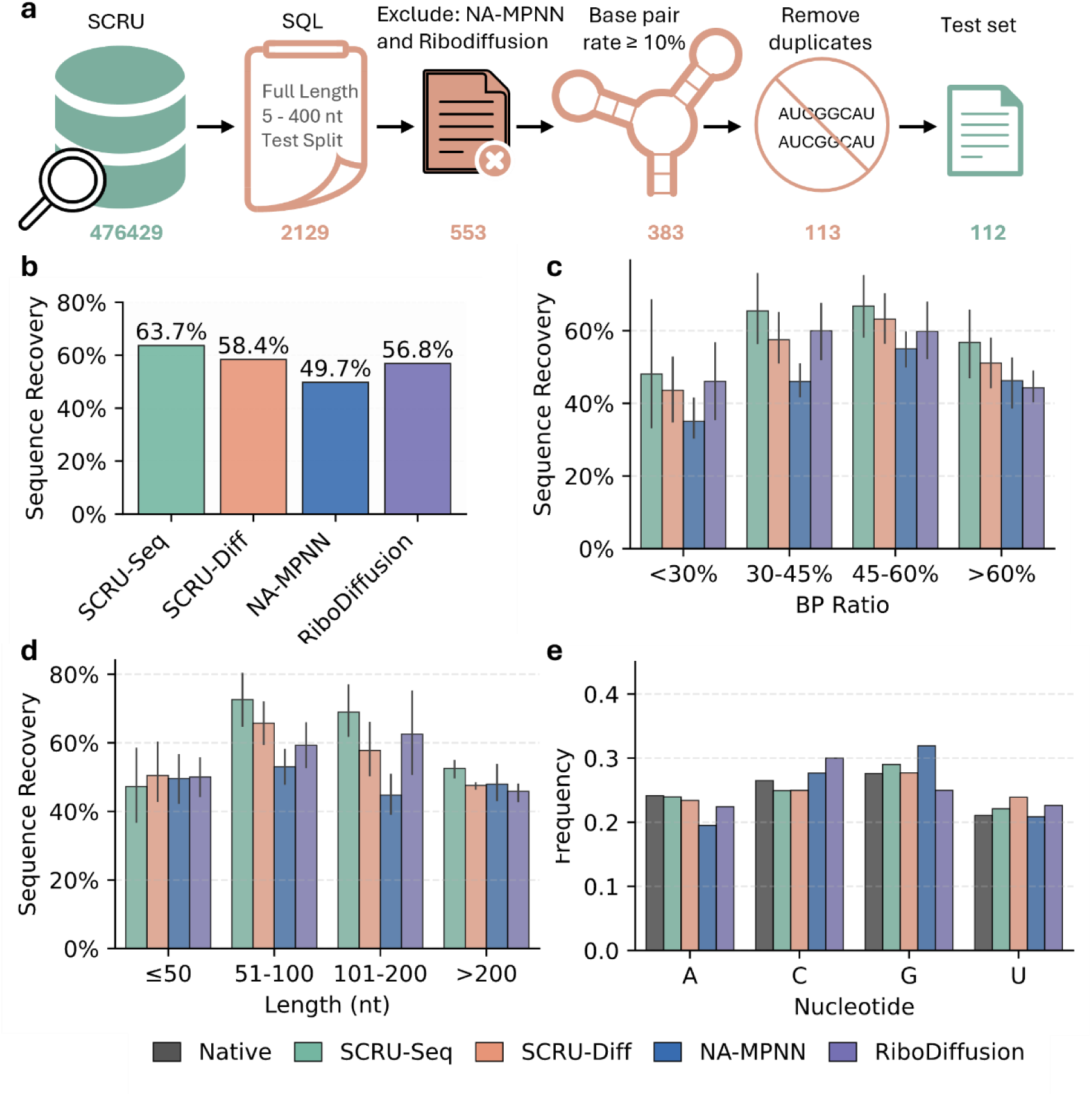
Sequence Recovery Benchmarking. Evaluation of Native Sequence Recovery (NSR)—the percentage of nucleotides identifying correctly with the original crystal structure—on the 112-candidate test set (set112). - a: Data filtering workflow to curate the non-redundant independent test set. - b: Overall NSR comparison across models, showing SCRU-Seq achieving 63.7%, significantly outperforming NA-MPNN (49.7%) and RiboDiffusion (56.8%). - c, d: Performance breakdown by Base-Pairing (BP) Ratio (c) and chain length in nucleotides (d). - e: Nucleotide frequency distribution. Gray bars represent the native PDB distribution, showing that SCRU-based models successfully capture the natural sequence composition.

To ensure parity in benchmarking against external baselines, we strictly excluded any PDB entries that appeared in the training sets of NA-MPNN and RiboDiffusion, reducing the pool to 553 candidates. After further filtering for secondary structure quality (≥10% base-pairing), removing redundant sequences, and enforcing 100% backbone atom completeness, we arrived at a final high-fidelity benchmark of 112 diverse RNA candidates. This strict curation guarantees that the evaluation represents a truly “double-blind” test for all models involved.

We specifically examined performance across two critical structural axes: base-pairing (BP) ratio and nucleotide length. The BP ratio serves as a proxy for structural density and complexity; while high-BP regions (stems) are governed by canonical constraints, low-BP regions (loops and junctions) often involve complex non-canonical interactions that represent a significant challenge for generative models. By tracking NSR against BP density (Figure 3c), we confirm that our models maintain high-fidelity design performance regardless of local structure type. Similarly, because long RNA chains introduce significant long-range dependencies and increase the potential for cumulative inference errors, we evaluated scalability across varying sequence lengths (Figure 3d). We find that both models maintain consistent accuracy from 20 to 400 nt, demonstrating that our modular training strategy on SCRU-DB effectively generalizes across diverse biological scales.

Notably, we show that the performance gap between the two parallel models is narrow across these dimensions. We further demonstrate that both models generate sequences with a nucleotide frequency distribution that aligns closely with biological native compositions (Figure 3e), confirming that our modeling strategies accurately capture the underlying structural constraints without over-optimizing toward specific sequences.

### Evaluation of 3D Structural Fidelity

Beyond sequence recovery, we assessed whether our designed sequences can actually adopt the target 3D backbone geometry. However, because accurate 3D structure prediction remains a significant challenge for genomic RNA, we first established a high-confidence benchmark for our evaluation tool. We performed “forward folding” on all 112 native RNA sequences from our test set using Boltz-1 and measured the deviation from their crystal structures. To ensure that our assessment of redesigned sequences was not confounded by the limitations of the prediction model, we selected a subset of 13 representative RNA targets where Boltz-1 demonstrated superior predictive accuracy for the native sequence (Figure 4a). This rigorous selection ensures that the structural metrics we report are grounded in Targets where the folding landscape is correctly captured by the evaluation engine.

**Figure 4:**
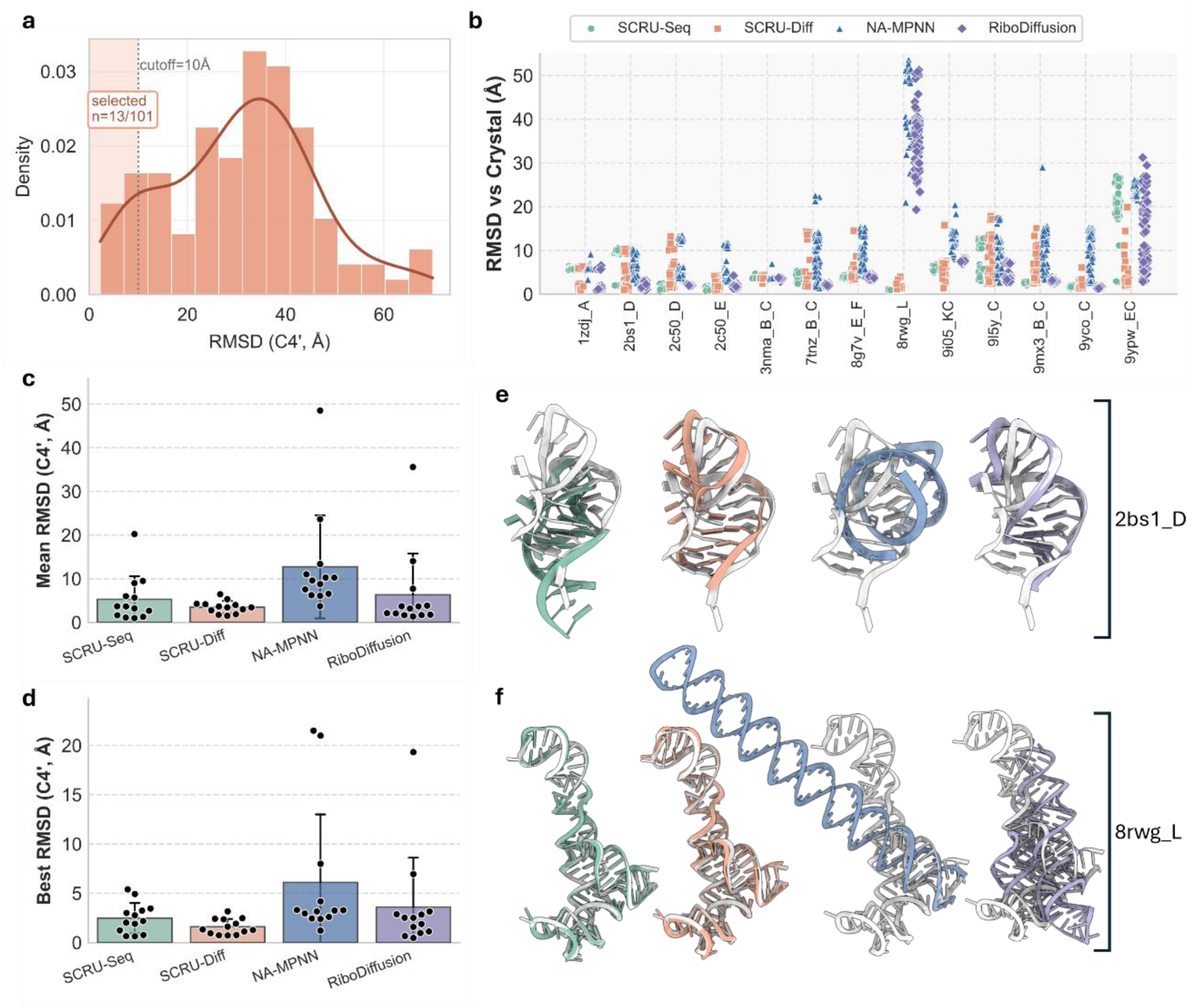
Structural Fidelity and RMSD Evaluation. Assessment of folding accuracy for designed sequences using C4’ RMSD (Root Mean Square Deviation focus on the C4’ ribose atom as a stable backbone anchor). - a: Density distribution of C4’ RMSD for candidates selected from the benchmark. High-fidelity targets (RMSD < 10Å) were identified for detailed 3D evaluation. - b: Individual RMSD values against experimental crystal structures across 13 diverse benchmarks. - c, d: Comparison of Mean RMSD (average across 100 samples) and Best RMSD (the highest accuracy solution found). SCRU-Diff achieves a Best RMSD of approximately 1.5Å for multiple targets. - e, f: Visual superposition of native crystal structures (gray) with designs from SCRU-Seq (teal), SCRU-Diff (salmon), NA-MPNN (blue), and RiboDiffusion (purple) for targets 2BS1_D (e) and 8RWG_L (f).

On this high-fidelity subset, we quantified structural fidelity by measuring the Root-Mean-Square Deviation (RMSD) between the 3D structures predicted for our designs and their native crystal structures (Figure 4). Specifically, we report the C4’ RMSD, utilizing the C4’ atom as the primary backbone anchor—the RNA equivalent to the C-alpha atom in proteins. Because the C4’ atom occupies a central position within the ribose sugar and maintains a relatively constant virtual bond distance (∼5.9 Å) between consecutive residues, it provides a clean, continuous trace of the global RNA trajectory. Unlike more flexible phosphate groups or side-chain-dependent base atoms, the C4’ trace captures the core topological fold while remaining robust to local thermal fluctuations in structural data.

We observe that the majority of our designs exhibit tight structural clustering with RMSD values below 10Å (Figure 4a). Across the 13 representative benchmarks, both SCRU-Seq and SCRU-Diff maintain high structural fidelity (Figure 4b). Crucially, we evaluated both the Mean RMSD (representing average predictive reliability) and the Best RMSD (the highest capability of the model’s generative ensemble). While both models are competitive in Mean metrics (Figure 4c), SCRU-Diff achieves a significantly lower Best RMSD, reaching approximately 1.5Å for multiple targets (Figure 4d). This gap between Mean and Best suggests that while direct prediction provides a stable baseline, the stochastic nature of the diffusion process allows SCRU-Diff to more effectively explore the structural landscape and identify high-fidelity conformational solutions.

Through visual superposition (Figure 4e, f), we demonstrate that designs for complex targets—specifically the large ribosomal fragment (PDB ID: 4V4G_W) and the multi-way junction (PDB ID: 8BU8_A)—successfully reproduce the native backbone architecture.

Our models correctly position branching hubs and long peripheral helices, maintaining the global topology essential for biological function. These results indicate that our modular training approach enables the generation of sequences that are structurally viable at various scales. This exceptional “Best Case” performance by SCRU-Diff directly motivates a deeper examination of its generative diversity in the following section.

### Generative Diversity and Exploration of Sequence Space

A primary objective in RNA design is to capture the “one-to-many” nature of the sequence-structure relationship: for any given 3D fold, a vast landscape of sequence solutions often exists. Evaluating the generative diversity of our models is critical because high-diversity sequence sets provide more candidates for downstream experimental optimization, allow for the balancing of multiple therapeutic constraints (e.g., GC content, stability, or lack of off-target motifs), and demonstrate that the model has learned the underlying structural rules rather than simply memorizing specific sequence templates. We therefore investigated the extent to which our models explore this generative space while maintaining structural validity (Figure 5).

**Figure 5:**
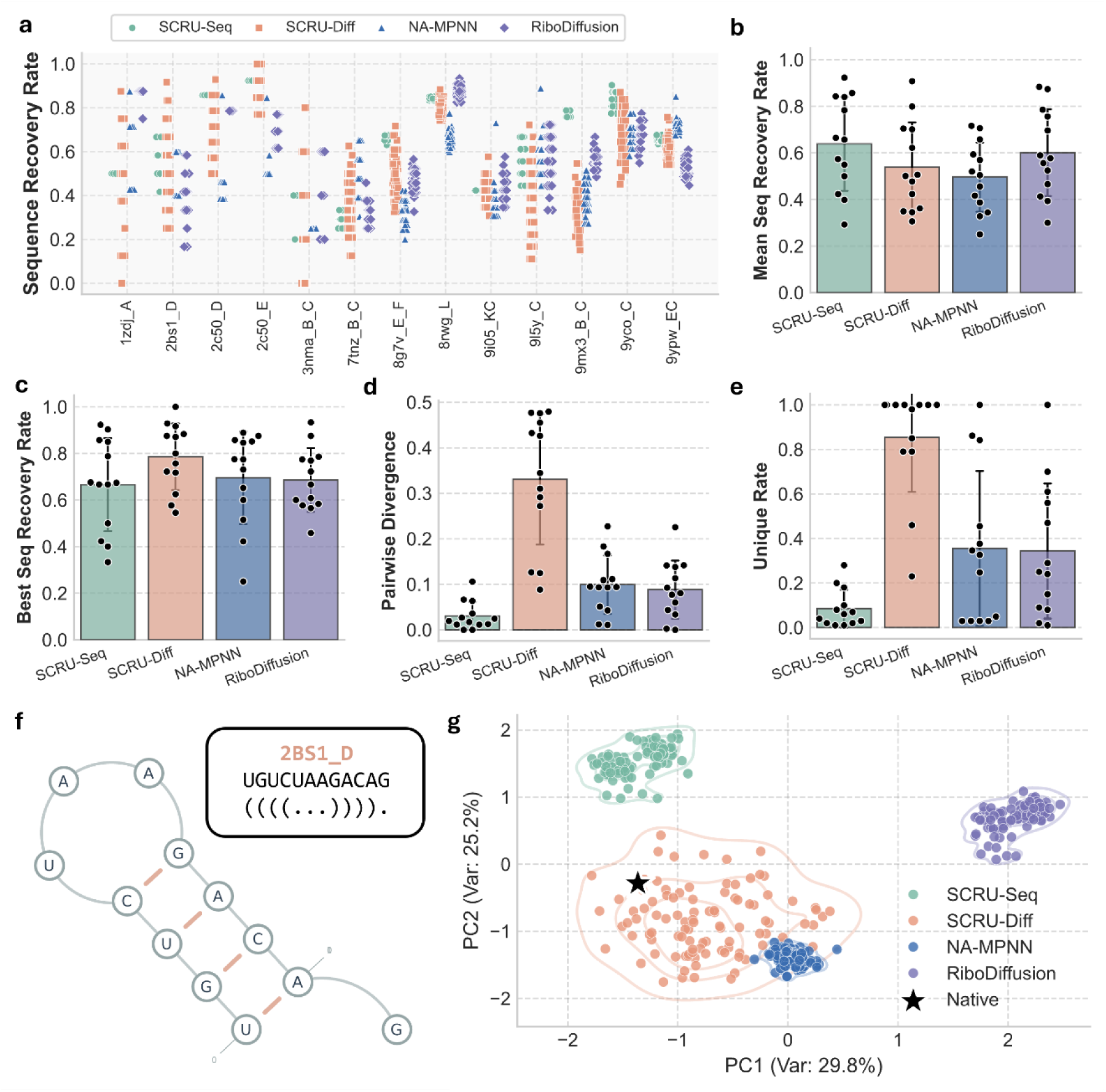
Sequence Diversity and Generative Space. Analysis of the sequence variations and exploration efficiency of different modeling paradigms. - a: Sequence recovery rate per target across the benchmark. - b, c: Comparison of Mean NSR (b) and Best NSR (c). SCRU-Diff reaches a superior Best NSR of 79.2%. - d, e: Pairwise Sequence Divergence (d) and Unique Sequence Rate (e). These metrics quantify generative diversity, showing SCRU-Diff’s ability to explore a much broader sequence landscape than deterministic or autoregressive baselines. - f: Secondary structure diagram of target 2BS1_D used as an example for diversity analysis. - g: Principal Component Analysis (PCA) of the sequence space. The distribution of SCRU-Diff designs (salmon) overlaps significantly with the native cluster (black star), demonstrating effective coverage of the functional sequence space.

We identify a core strength of our SCRU-based framework, particularly the iterative SCRU-Diff model, in its ability to navigate this broad space. We find that while SCRU-Seq provide the highest average predictive accuracy, SCRU-Diff achieves a superior Best NSR of 79.2% (Figure 5c) across 100 samples per target, notably exceeding the best-case recovery of RiboDiffusion (67.4%) and NA-MPNN (58.1%). More importantly, we demonstrate that SCRU-Diff produces significantly higher Unique Sequence Rates (∼85%) and Pairwise Divergence (∼0.33) than both the direct-prediction model and external baselines (Figure 5d, e). While RiboDiffusion also employs a diffusion paradigm, our structural decomposition on SCRU-DB allows SCRU-Diff to generate more distinct and structurally viable sequence variations. Our Principal Component Analysis (PCA) of the sequence space (Figure 5g) quantitatively confirms this: the SCRU-Diff distribution is significantly broader and successfully encompasses the native sequence cluster (black star), whereas NA-MPNN, RiboDiffusion, and SCRU-Seq often produce more restricted, tightly clustered designs that fail to capture the full breadth of the biological solution space.

### Structural Stability and Context Independence of SCRUs

A fundamental assumption of our framework is that the modular SCRU units are physically self-sufficient—that they maintain their structural motifs even when extracted from their global RNA context. To validate this assumption, we performed a computational study to assess the structural isomorphism of SCRUs: whether their local fold in isolation matches their role in the complete RNA chain (Figure 6). Because most SCRUs are fragments of larger RNAs, we must confirm that their local sequence is sufficient to encode the target geometry without distal stabilization. We utilized secondary structure (SS) folding as a proxy, comparing the predicted folds for SCRUs across three states: Isolation (ISO), where the designed SCRU sequence is folded alone; Native (NAT), representing the original crystal structure environment; and Contextual (CTX), where the designed SCRU sequence is embedded within the full-length redesigned RNA chain (Figure 6a). This triple-comparison allows us to determine if the surrounding redesigned context destabilizes or alters the intended local fold. Notably, to ensure a reliable baseline, we focused our analysis on targets where the native sequence’s crystal structure fold was accurately recovered by the prediction model (UFold).

**Figure 6:**
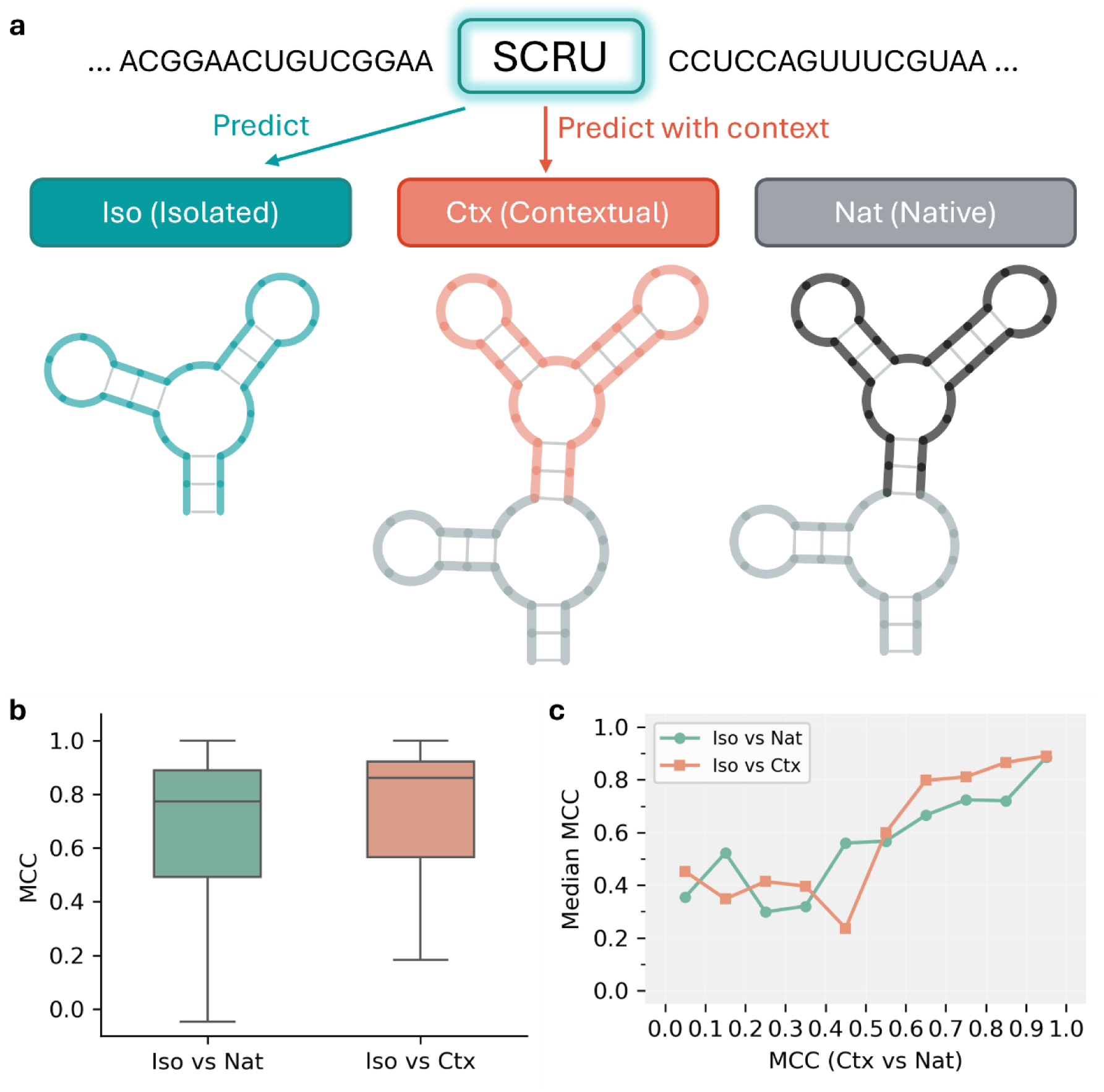
Evaluation of Structural Stability and Context Independence. Assessing the structural isomorphism and self-sufficiency of modular SCRU units. - a: Schematic comparing fold predictions across three environments: Isolation (ISO) (designed SCRU folded alone), Native (NAT) (native crystal structure environment), and Contextual (CTX) (designed SCRU embedded in the full-length redesigned chain). - b: Matthews Correlation Coefficient (MCC) distribution. MCC is used here as a balanced measure of secondary structure folding fidelity. The high median MCC (0.86) for Iso vs. Ctx confirms that SCRUs act as independent structural modules. - c: Relationship between median MCC and context dependence (defined by MCC of Ctx vs. Nat), demonstrating that SCRUs remain thermodynamically stable regardless of their global sequence environment.

Our Matthews Correlation Coefficient (MCC) analysis shows that the majority of SCRUs act as thermodynamically stable and structurally consistent building blocks. We chose MCC as our primary metric because it provides a balanced evaluation of base-pairing fidelity, effectively accounting for the inherent class imbalance between paired and unpaired nucleotides. Unlike simple accuracy, MCC penalizes both false positives and false negatives, ensuring that the high scores reflect true structural isomorphism rather than over-prediction of either state. We find that the median MCC for Isolated vs. Contextual reached 0.86, significantly higher than the Isolated vs. Native baseline (0.78) (Figure 6b). The high level of agreement between these states—where the isolated SCRU fold precisely recapitulates its contextual structural role (Figure 6c)—confirms that SCRUs act as independent structural modules. We thus validate our fundamental hypothesis that training on modular, context-independent fragments—enabled by the SCRU-DB—provides a cleaner and more physically grounded sequence-structure mapping than traditional global modeling approaches.

## Discussion

The central tenet of our work is that RNA structures can be split into SCRUs that exhibit high structural stability independent of their global environment. By demonstrating that isolated SCRUs consistently maintain their native dot-bracket folds (as validated by UFold benchmarks), we provide empirical evidence that the local sequence-structure mapping in RNA is exceptionally robust. This modularity is the key to our framework’s performance: by training on these diverse, self-contained units from SCRU-DB, our models learn the fundamental “grammar” of RNA folding that is transferable across different biological contexts.

A critical contribution of this work lies in the architectural innovations of SCRU-Seq, specifically designed to overcome the limitations of traditional graph representations for RNA. Standard GNN approaches often struggle to balance local chemical fidelity with global structural context. A pure k-nearest neighbor graph can miss long-range tertiary contacts essential for folding, while a fully connected graph becomes computationally prohibitive. To resolve this, SCRU-Seq introduces a Dual-Radius Graph architecture that operates at two complementary scales. At the atomic scale, dense all-atom edges within a 4Å radius capture precise stereochemical constraints such as sugar pucker configurations, phosphate backbone torsion angles, and non-canonical hydrogen bonding networks, ensuring that the generated sequence respects the fundamental physicochemical laws of RNA stability. Simultaneously, at the structural scale, the model maintains a sparse, long-range graph connecting C4’ atoms up to 20Å, effectively “short-circuiting” the graph to allow information to propagate rapidly across the entire molecule. This global view is crucial for resolving the sequence identity of distal interacting loops or stems that, while chemically distant in sequence space, are spatially adjacent in the 3D fold.

In deep Graph Neural Networks, a common failure mode is “over-smoothing,” where node representations become indistinguishable after many layers of message passing. This is particularly detrimental for RNA design, where a single nucleotide mutation can drastically alter the folding landscape. To combat this, SCRU-Seq employs a Gated Message Passing mechanism. By modulating the information flow with learnable gates (sigmoid activation), the network can selectively filter noise while amplifying critical structural signals. This allows us to train a deeper network (16 layers) without signal degradation, ensuring that the final node embeddings retain distinct, high-resolution information about their local structural role.

The prevailing assumption in sequence design has been that sophisticated inference strategies—Autoregressive (AR) generation or Diffusion-based sampling—are necessary to capture the complex inter-dependencies between residues. However, our results with SCRU-Seq challenge this assumption at its foundation. We argue that the reliance on AR and Diffusion is fundamentally a compensatory mechanism for data scarcity. When trained on limited PDB data, these complex inference procedures squeeze marginal performance from insufficient training examples. NA-MPNN employs autoregressive decoding, generating sequences one nucleotide at a time, while RiboDiffusion requires hundreds of iterative denoising steps. Both methods add substantial computational overhead to compensate for what is essentially a data limitation.

Our research demonstrates that state-of-the-art performance—63.7% native sequence recovery for our models compared to 49.7% for NA-MPNN and 56.8% for RiboDiffusion—can be achieved without relying on expensive global Diffusion alone. Instead, we propose two parallel strategies: SCRU-Seq, a high-speed direct prediction baseline, and SCRU-Diff, a generative diffusion model. Both models leverage an expanded training dataset of 476,429 units from 8,888 PDB entries (a 53-fold expansion) and a multi-scale graph representation. The Dual-Radius Graph architecture—capturing both atomic-scale chemistry and global topology—effectively “pre-computes” the dependencies that AR models learn sequentially. By evaluating SCRU-Seq as a direct prediction point, we highlight that even simpler models can achieve high accuracy when trained on SCRU-DB. However, SCRU-Diff further pushes the boundaries of design by providing high diversity and competitive accuracy, reaching a Best NSR of 79.2%. This iterative model excels at capturing the one-to-many nature of the RNA design problem, effectively exploring the sequence landscape to identify highly accurate solutions—validated by C4’ RMSD values as low as 1.5Å for multiple complex targets. This validates our core hypothesis: the bottleneck in RNA design is data accessibility and granularity, not model complexity. With the right modular representation (SCRU-DB), both direct and iterative models can efficiently solve the inverse folding problem.

Furthermore, our validation of SCRU modularity highlights a critical methodological choice in assessing RNA design. While the “gold standard” for fragment stability would ideally involve 3D geometric comparisons (e.g., C4’ RMSD), current 3D structure prediction models—including state-of-the-art tools like Boltz-1—remain substantially less accurate when applied to small, diverse RNA fragments compared to full-length genomic RNAs. This inaccuracy introduces significant noise that can confound structural assessments. Consequently, we employed 2D secondary structure metrics as a robust surrogate to evaluate context-independence and thermodynamic viability. By demonstrating that isolated SCRU sequences reliably recover their target dot-bracket motifs—thereby establishing structural isomorphism between the fragment and its parent RNA—we confirm that our decomposition strategy identifies physically self-stabilizing units that can serve as reliable building blocks for scalable RNA design.

Several limitations remain. First, our current framework does not explicitly model tertiary interaction constraints such as long-range base triples and non-canonical stacking interactions that contribute to RNA stability. Second, extension to DNA-RNA and Protein-RNA complexes would broaden the applicability of our approach but requires additional architectural adaptations. Third, experimental validation of designed sequences through in vitro assays remains essential to confirm the folding and functional properties predicted by the model. Future work will address these challenges while maintaining the data-centric philosophy that underlies our approach.

## Methods

### SCRU-DB: Modernized RNA Structural Motif Engineering

To address the critical limitations of existing RNA datasets characterized by inconsistent quality control, limited scale, and inadequate structural decomposition, we developed the SCRU-DB (Self-contained RNA Unit Database). This resource transforms the raw RCSB Protein Data Bank landscape into a high-fidelity, searchable structural database optimized for deep learning applications. Existing RNA structure datasets suffer from several fundamental shortcomings: they often include low-resolution or incomplete structures, lack systematic decomposition into functional units, and provide insufficient annotation of molecular interactions. SCRU-DB addresses these issues through a comprehensive ETL pipeline with rigorous quality control, a hybrid database architecture for efficient data access, and a hierarchical decomposition algorithm that identifies 61,916 structurally meaningful RNA units (SCRUs).

ETL Pipeline and Hybrid Database Architecture. The pipeline processes approximately 170,000 RNA chains from the PDB, extracting structural information from mmCIF format files. We utilize an ETL (Extract-Transform-Load) architecture designed for scalability, where each PDB entry undergoes a series of transformations: structural parsing, residue renumbering, secondary structure detection, SCRU extraction, and database storage. Structural mmCIF files are parsed using OpenStructure (OST), a C++-based molecular structure library, extracting all 12 backbone atoms (P, OP1, OP2, O5’, C5’, C4’, O4’, C3’, O3’, C2’, O2’, C1’) for each nucleotide. To balance relational querying efficiency with heavy coordinate storage, we implemented a hybrid database system that leverages the strengths of both systems. DuckDB serves as the metadata index, supporting rapid relational joins on sequences, secondary structures, and interaction labels with vectorized query execution. Sharded SQLite stores all-atom coordinates as binary-serialized dictionaries, partitioned across multiple files to ensure O(1) random access of 3D geometry during multi-worker training. This architecture enables efficient parallel data loading while maintaining flexible querying capabilities. The sharding strategy distributes data across approximately 100 SQLite files, each containing roughly 1,000-2,000 RNA chains, enabling balanced load distribution across worker processes.

Physical Base-Pairing Recognition Engine. Accurate identification of canonical base pairs is fundamental to RNA structure analysis. We implemented a high-performance Python-based pairing engine that identifies canonical Watson-Crick (A-U, G-C) and Wobble (G-U) pairs by evaluating 3D spatial constraints and hydrogen-bond (H-bond) energetics without external dependencies. The engine calculates a cumulative probability score, 𝑆_𝑡𝑜𝑡𝑎𝑙_ = ∑_𝑖_ 𝑠_𝑖_, where for each potential donor-acceptor pair, the score 𝑠_𝑖_ is derived from a Gaussian decay function centered at the optimal H-bond distance: 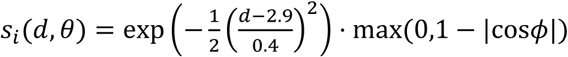. This formulation captures both distance dependence (favoring ∼2.9 Å separations) and angular dependence (favoring perpendicular H-bond orientations). Pairs are recognized as canonical when 𝑆_𝑡𝑜𝑡𝑎𝑙_ > 0.8, a threshold determined empirically through validation against annotated base pairs in the PDB. To filter out non-canonical interactions that may pass the H-bond score threshold, we enforce strict geometric constraints: base ring normals must be anti-parallel (n_𝑖_ ⋅n_𝑗_ < −0.5) to ensure coplanar stacking, and the glycosidic configuration must be in the Cis configuration (|𝜒| < 𝜋/2) characteristic of standard RNA helices. This pairing engine enables systematic secondary structure annotation across the entire PDB without manual curation.

Graph-based Structural Partitioning of SCRUs. Central to our framework is the extraction of SCRUs using a graph-based partitioning algorithm that naturally accommodates complex topologies, including pseudoknots. Unlike traditional motif extraction that isolates single loops or relies on a nested tree hierarchy, our algorithm identifies autonomous structural modules by treating helical regions (stems) as fundamental nodes of stability—owing to the dense network of canonical base pairs—and the intervening loops or junctions as linking edges. The partitioning process begins by mapping an RNA chain into a connectivity graph where nodes are helical fragments and edges represent sequence segments navigating between them. These elements are then selectively combined into "self-contained" clusters through a weight-constrained greedy growth protocol. A cluster expands from a seed edge by incorporating adjacent helices and fragments as long as they are linked by strong tertiary interactions (defined by at least five non-canonical base pairs) or short helical bridges (ten residues or fewer) that fail to insulate the motifs from each other. This graph-based strategy identifies robust, physically contiguous clusters where the local fold is stabilized by its internal pairing network. This definition ensures that each SCRU serves as a chemically complete and structurally isomorphic prior—retaining its precise native fold regardless of its global environment—for high-fidelity sequence design..

Interaction Annotation. Real-world RNA molecules rarely exist in isolation; they engage in diverse molecular interactions with proteins, DNA, and small molecule ligands that can substantially influence their structure and function. For every RNA residue, we perform a 4.0 Å spatial search via KDTree to detect interactions with proteins, DNA, and small molecule ligands. This distance threshold captures both direct hydrogen bonding and van der Waals contacts that may influence local structure. The interaction information is encoded as categorical labels (‘P’ for protein contact, ‘D’ for DNA contact, ‘L’ for ligand contact, ‘.’ for unconstrained), enabling the model to learn context-dependent sequence preferences at molecular interfaces. This annotation is particularly valuable for designing RNA interfaces in therapeutic applications, such as aptamer development or RNA-protein interaction engineering.

### SCRU-Seq: Direct-Prediction Generative Model

SCRU-Seq is a Graph Neural Network (GNN) model optimized for high-speed RNA design through a high-efficiency direct-prediction paradigm. The inverse folding problem—the task of designing an RNA sequence that will fold into a given target structure—has emerged as a central challenge in RNA bioinformatics and synthetic biology. SCRU-Seq addresses this through an innovative dual-radius graph representation and a non-autoregressive architecture that predicts sequences in a single forward pass.

Dual-Radius Graph Representation. Instead of a simple k-nearest neighbor graph that treats all connections equally, we represent the RNA backbone using a Dual-Radius Relational Graph that captures both local chemical environments and global topological relationships at distinct scales. At the atomic scale, edges connect all 12 backbone atoms within 4 Å to capture steric constraints, local bonding patterns, and chemical environment. This radius is sufficient to capture covalent connectivity, base-stacking interactions, and immediate steric clashes. At the structural scale, edges connect C4’ atoms within 20 Å to capture the global folding topology without saturating the graph with excessive edges. The choice of C4’ as the structural anchor reflects its central position in the ribose sugar and its consistent availability across all nucleotides. This dual-radius design balances computational efficiency with structural expressivity: the atomic radius ensures the model has access to sufficient local chemical information, while the structural radius provides long-range connectivity that informs global fold topology. Each atom node is featurized with a 12-dimensional one-hot encoding for atom type (distinguishing P, OP1, OP2, O5’, C5’, C4’, O4’, C3’, O3’, C2’, O2’, C1’), a 32-dimensional sinusoidal positional encoding to preserve sequential order along the RNA chain, and semantic masks indicating prediction targets and interaction status.

Model Architecture. The model core consists of 16 Graph Convolutional Layers with bidirectional message passing, enabling information to flow in both directions along the graph edges. Each layer employs a multiplicative gating mechanism (ℎ_𝑚𝑠𝑔_ ⋅ 𝜎(ℎ_𝑔𝑎𝑡𝑒_)) that dynamically controls information flow and prevents gradient vanishing during deep propagation. The gating mechanism acts as a learned attention filter, allowing the model to selectively propagate relevant information while suppressing noise. Information is updated symmetrically across edges, combining source and destination node features with edge attributes including Euclidean distance (representing spatial separation) and edge type (distinguishing atomic-scale vs. structural-scale connections). We utilize SiLU (Swish) activation and LayerNorm for training stability, following best practices from modern deep learning architectures. The output layer projects node features to a 4-class nucleotide distribution (A, U, G, C) at each C4’ position, where supervision is applied during training.

Direct Prediction Paradigm. Unlike traditional autoregressive models that generate sequences residue-by-residue, requiring sequential sampling and suffering from error accumulation, SCRU-Seq utilizes a Direct Prediction (One-Shot) architecture. The model is trained to map the 3D backbone geometry directly to the probability distribution of all nucleotides simultaneously, eliminating exposure bias and sequential error accumulation. Because the entire sequence is predicted in a single forward pass, the model achieves orders-of-magnitude faster inference compared to baseline methods (∼100x faster than NA-MPNN), making it ideal for large-scale library generation.

### SCRU-Diff: Iterative Diffusion Generative Model

To provide a high-diversity alternative and explore the sequence space more thoroughly, we developed SCRU-Diff, a generative diffusion model that operates in parallel with the direct prediction of SCRU-Seq. SCRU-Diff is designed to generate highly diverse and structurally optimized sequences for a single target backbone through a stochastic denoising process.

Discrete Diffusion Probabilistic Model (D3PM). SCRU-Diff utilizes a Discrete Diffusion (D3PM) framework acting on the four-letter nucleotide alphabet (𝛴 = {𝐴, 𝑈, 𝐺, 𝐶}). During the forward process, noise is added to the native sequence via a transition matrix 𝑄_𝑡_, eventually reaching a uniform stationary distribution. The model is trained to reverse this process, learning to denoise a random sequence back into a structurally valid candidate. We employ a linear noise schedule typically spanning 100 to 1,000 steps, allowing for fine-grained control over the generative trajectory.

Iterative Refinement and Diversity. By leveraging the same Dual-Radius Graph architecture as SCRU-Seq, SCRU-Diff captures long-range structural dependencies during each denoising step. Unlike the deterministic nature of direct prediction, SCRU-Diff’s stochastic sampling allows it to explore multiple distinct regions of the sequence-structure landscape. This results in a significantly higher Unique Sequence Rate and Pairwise Divergence, as the model can generate a broad ensemble of candidates that all satisfy the target backbone constraints.

### Training Protocol and Data Integrity

Dataset Configuration. Models are trained on the SCRU-DB dataset using a configurable dataset system that supports diverse RNA modeling tasks. The CoarseDataset class provides C4’ coordinate-based training with one-hot nucleotide features, representing RNA at the backbone level suitable for sequence design. The system supports configurable maximum sequence length (400 nt), positional encoding dimension (32), and batch collation with automatic padding for variable-length sequences. A three-tier caching system (memory/file/none) enables optimization for different computational setups: memory caching for single-GPU training where dataset size fits in RAM, file-based caching for distributed training across multiple nodes, and disable mode for debugging or streaming very large datasets. The dataset configuration is managed through a central DATASET_CONFIGS dictionary that specifies dataset class, feature dimensions, and processing parameters for each task.

Optimization Strategy. Training uses AdamW optimizer with learning rate 1e-4, weight decay 1e-12, and amsgrad enabled for improved convergence. Models are trained with batch size 8 (distributed across GPUs) for 500 epochs, with early stopping based on validation loss plateau. The training framework is built on PyTorch Lightning, providing automated checkpointing, gradient accumulation, and mixed-precision training (FP16) for memory efficiency and speed. Validation metrics are aggregated epoch-wise using custom aggregation functions, with detailed logging of training progress including loss curves, accuracy metrics, and timing statistics.

Data Integrity and Split Strategy. Preventing data leakage between training and test sets is critical. To achieve this, we perform dataset splitting at the cluster level. All unique sequences in SCRU-DB are clustered using MMseqs2 with 80% identity and coverage thresholds, identifying approximately 8,405 non-redundant sequence clusters (including sub-fragments and full-chains). A deterministic hash function assigns clusters to train/val/test splits with an 80/10/10 ratio. All SCRUs belonging to the same cluster are assigned to the same split, and motifs from the same PDB entry are kept within the same split to avoid intra-structure leakage. Quality filtering retains only RNA chains with 100% backbone atom completeness, resolution ≤ 3.5 Å, and at least 10% standard base-pairing.

### Benchmarking and Evaluation

Test Set Selection. We curated an independent benchmark of full-length RNA chains to ensure a rigorous and unbiased evaluation through a strict sequential filtering pipeline (Figure 3a). Starting with an initial pool of 2,129 full-chain SCRUs assigned to the test split (5–400 nt), we first applied a baseline exclusion step to prevent data leakage, removing all PDB entries present in the training sets of NA-MPNN and RiboDiffusion and leaving 553 candidates. To ensure each candidate provided a well-defined structural template, we required at least 10% standard Watson-Crick or Wobble base pairing, reducing the pool to 383 RNAs. We then performed sequence deduplication to ensure a non-redundant set of structural challenges, resulting in 113 candidates. Finally, we enforced a strict requirement for 100% backbone atom completeness (at least 12 heavy atoms per residue), yielding a final high-fidelity benchmark of 112 non-redundant RNA chains.

This meticulous curation ensures that our evaluation set contains only high-quality, biologically relevant structural templates that are completely unseen by any of the compared models.

Evaluation Metrics. We employ two complementary metrics that capture different aspects of design quality. Native Sequence Recovery (NSR) measures the percentage of residues where the top-1 predicted nucleotide matches the native PDB sequence, providing a direct measure of how well the model recovers the evolutionary-optimized sequence. While high NSR indicates structural validity, we recognize that many sequences may fold into the same structure, motivating our second metric. 2D Folding F1 assesses structural fidelity by folding the designed sequence using UFold, a deep learning-based secondary structure predictor, and computing the F1 score of recovered base pairs relative to the native dot-bracket structure. High 2D F1 confirms that the designed sequence creates a stable fold matching the target geometry, even if the exact sequence differs from the native. The combination of NSR and 2D F1 provides a comprehensive evaluation: NSR measures sequence recovery while 2D F1 measures fold preservation.

Baseline Models. We compare against two established RNA design methods that represent the current state-of-the-art. NA-MPNN (Jia et al., 2023) uses autoregressive message passing with per-residue output, representing the leading autoregressive approach. RiboDiffusion (Khavnekar et al., 2023) employs continuous diffusion models for RNA structure generation, representing the emerging diffusion-based paradigm. All baseline models are evaluated using their recommended configurations on the same set50_all test set, ensuring methodological parity in evaluation. We report both NSR and 2D F1 for all methods to enable comprehensive comparison across different aspects of design quality.

Stability Validation. A unique contribution of our framework is the validation of SCRU structural stability—testing whether extracted units maintain their fold when isolated from the parent complex. This is essential for understanding whether SCRUs are truly independent structural modules or require context for proper folding. Using UFold for secondary structure prediction, we compute the Matthews Correlation Coefficient (MCC) between in-context predictions (folding the full RNA chain containing the SCRU) and in-isolation predictions (folding the SCRU alone). An MCC of 1.0 indicates complete self-reliance of the structural motif, meaning the SCRU contains all information needed to specify its fold. Lower MCC values indicate context dependence, valuable information for understanding RNA folding principles and informing design strategies.

### Computational Requirements

Training requires 8x A100 GPUs (40GB) for approximately 500 epochs, with total training time of approximately 1-2 weeks depending on dataset size and augmentation settings. Single-GPU inference is supported with reduced batch sizes, enabling deployment on more modest hardware. The hybrid database architecture ensures efficient data loading with O(1) random access to coordinate geometry, minimizing I/O bottlenecks during training. Memory requirements are dominated by model parameters (approximately 50-100M parameters for the full model) and batch storage for graph representations.

## ACKNOWLEDGMENTS

We thank the Research Computing at The University of Virginia for providing computational resources and technical support that have contributed to the results reported within this publication. URL: https://rc.virginia.edu.

## COMPETING INTERESTS

All authors declare no competing interests.

### Data, code, and materials availability

All data and code needed to evaluate and reproduce the conclusions in the paper are present in the paper and/or the Supplementary Materials. No new materials were generated in this study. Source codes will be released soon.

